# Fully automated and integrated proteomics sample preparation platform for high-throughput drug target identification

**DOI:** 10.1101/2023.07.14.548974

**Authors:** Qiong Wu, Jiangnan Zheng, Xintong Sui, Changying Fu, Xiaozhen Cui, Bin Liao, Hongchao Ji, Yang Luo, An He, Xue Lu, Chris Soon Heng Tan, Ruijun Tian

## Abstract

With the increased demand of large-cohort proteomic analysis, fast and reproducible sample preparation has become the critical issue that needs to be solved. Herein, we developed a fully automated and integrated proteomics sample preparation workflow (autoSISPROT), enabling the simultaneous processing of 96 samples in less than 2.5 hours. Benefiting from its 96-channel all-in-tip operation, protein digestion, peptide desalting, and TMT labeling could be achieved in a fully automated manner. The autoSISPROT demonstrated good sample preparation performances, including >94% of digestion efficiency, nearly 100% of alkylation efficiency, >98% of TMT labeling efficiency, and >0.9 of intra- and inter-batch Pearson correlation coefficients. Furthermore, by combining with cellular thermal shift assay-coupled to mass spectrometry (CETSA-MS), the autoSISPROT was able to process and TMT-label 40 samples automatically and accurately identify the known target of methotrexate. Importantly, taking advantage of the data independent acquisition and isothermal CETSA-MS, the autoSISPROT was well applied for identifying known targets and potential off-targets of 20 kinase inhibitors by automatedly processing 87 samples, affording over a 10-fold improvement in throughput when compared to classical CETSA-MS. Collectively, we developed a fully automated and integrated workflow for high-throughput proteomics sample preparation and drug target identification.

## Introduction

Mass spectrometry (MS)-based proteomics has emerged as the leading technology for systematically characterizing protein abundance, post-translational modifications, and protein-protein interactions, etc^1, 2^. For studying complex biological and clinical samples, large number of samples with multiple conditions and sufficient replicates are required to reach enough statistical power. However, sample preparation for high-throughput quantitative proteomic analysis remains a challenge^3^. Since traditional proteomics workflows typically require multi-step sample preparation, manually processing hundreds of samples is not only a very time-consuming and overwhelming task, but also introduces variation that affects the overall technical reproducibility. In order to standardize sample preparation, reduce time and labor costs, and improve reproducibility, there is a high demand for developing automated and high-throughput proteomics sample preparation methods.

In the past decade, various automated or high-throughput sample preparation methods have been developed that can be categorized into two types based on digestion mechanisms: in-solution digestion and on-bead digestion. For instance, Geyer et al. introduced a rapid and robust “plasma proteome profiling” pipeline according to in-StageTip (iST) method with an automated set-up on a liquid handling Bravo workstation (Agilent Technologies)^4, 5^. They applied this automated workflow for the in-depth study of 100 taxonomically diverse organisms^6^. To increase the analysis throughput of sample preparation, Messner et al. and Burns et al. reported ultra-high-throughput (384-well format) sample preparation platforms that implemented in-solution digestion strategy using the Biomek NXp and Bravo workstations, respectively^7, 8^. Although these methods demonstrated success in 96-well in-solution digestion, they are limited by long digestion time (typically overnight) and the need for multiple sample transfer steps^9^. More importantly, these methods require the off-line peptide desalting step, making it challenging to achieve fully automated sample preparation. By employing the on-bead digestion strategy, most of the proteomics sample preparation steps could be seamlessly integrated, thereby increasing sample preparation reproducibility. Building upon our previously developed simple and integrated spintip-based proteomics technology (SISPROT)^10, 11^, an on-bead digestion strategy, we have established a gas-pressure-driven microfluidic chip system for fully automated and integrated proteomics sample preparation in a single-channel manner^12^. Müller et al. implemented the single-pot solid-phase enhanced sample preparation (SP3) on a Bravo workstation (autoSP3) for automated processing of tissue lysates in a 96-well format^13, 14^. However, achieving fully automated sample preparation with autoSP3 was not user-friendly due to the requirement of off-line peptide desalting. It is worth noting that the aforementioned methods solely utilized the standard functions of Bravo workstation, which involve the aspiration and dispensing of liquids using disposable 96-well tips. However, the AssayMAP Bravo mode, a 96-well microchromatography platform based on disposable cartridges with packed bed of resins^15^, has not yet been investigated for fully automated proteomic sample preparation^16^. To date, a fully automated and high-throughput proteomic sample preparation workflow is still lacking.

The cellular thermal shift assay-coupled to mass spectrometry (CETSA-MS), also known as thermal proteome profiling (TPP), has emerged as a popular method to identify targets and off-targets of drugs based on ligand-induced changes in protein thermal stability^17–23^. Current implementations of CETSA-MS involve performing measurements at ten temperature points with two replicates per condition to estimate the thermal melting temperature (T_m_) shift between drug and vehicle treatment conditions. As a result, 40 samples need to be prepared and labeled with four sets of tandem mass tags (TMT), followed by off-line fractionation steps, which limits its throughput. To increase the analysis throughput of CETSA-MS, Gaetani et al. developed the proteome integral solubility alteration (PISA) by pooling the soluble fractions of multiple samples heated across a temperature gradient, resulting in a 10-20 times increase in throughput compared to CETSA-MS^24, 25^. Isothermal shift assay (iTSA) quantifies the difference in the soluble protein fraction at a single temperature, offering a 4-fold improvement in throughput compared to CETSA-MS^26^. However, both PISA and iTSA methods were limited to the labeling channel of TMT reagents, making it difficult to meet the demand for high-throughput drug target identification. Ruan et al. reported the matrix thermal shift assay (mTSA) by incorporating data independent acquisition (DIA) into iTSA, which improved throughput and was not limited to the TMT channel numbers^27^. Overall, for large-scale drug target identification via CETSA-MS, an automated and high-throughput sample preparation method is urgently needed.

In this study, we developed a fully automated and integrated 96-channel proteomic sample preparation platform named autoSISPROT. This platform combines the all-in-tip sample preparation capabilities of SISPROT with the precise liquid handling provided by AssayMAP Bravo^15^, enabling the simultaneous processing of 96 samples in 2.5 hours in a fully automated manner. We evaluated the sample preparation performance of autoSISPROT, as well as the intra- and inter-batch reproducibility, by processing a total of three 96-well plates on three different days. Furthermore, by combining with CETSA-MS, autoSISPROT can automatically process and TMT-label 40 samples and accurately identify the known target of the well-characterized model drug, methotrexate (MTX). Furthermore, we conducted a comprehensive assessment of two quantitative proteomic methods, namely TMT and DIA, for drug target identification by utilizing a pan-kinase inhibitor, staurosporine. Finally, to improve the analysis throughput of CETSA-MS, autoSISPROT and DIA-based CETSA-MS are combined to identify the known targets and potential off-targets of 20 kinase inhibitors in a fully automated and high-throughput manner.

## Results

### Development of a fully automated and integrated 96-channel proteomic sample preparation platform (autoSISPROT)

SISPROT allows for full integration of sample loading, protein reduction, alkylation, digestion, TMT labeling (optional), and desalting, all within a single spintip packed sequentially with C18 membrane and mixed SCX/SAX beads^10, 11, 28^. We hypothesized that this all-in-tip sample preparation could be automated on AssayMAP Bravo, a microchromatography platform with a 96-channel liquid handling head. This platform can precisely control the flow rates of upward aspiration and downward dispensing within the range of 2.5-20 μL/min, which allows us to program each sample preparation step on SISPROT-based cartridge with desired flow rate and processing time (**Fig. 1a**). Accordingly, we systematically optimized and significantly reduced the number of the buffers used in each step of sample preparation which allows the fully automated operation on the Bravo system with only 7 working decks (**Fig. 1b**). To ensure an ultra-low dead volume in autoSISPROT, we designed and fabricated the SISPROT-based cartridges that were assembled with the top and bottom tips with specialized shapes (**Fig. 1c**). The bottom component was packed with C18 membrane and mixed SCX/SAX beads in tandem. The preparation of SISPROT-based cartridges followed a strict ten-step process, and a three-step operating procedure was used as a rigorous quality control to ensure the reproducibility of the SISPROT-based cartridges. In summary, we designed the autoSISPROT protocol to automated process up to 96 samples in parallel within 2.5 hours, eliminating the need for manual intervention once the start button is clicked.

**Fig. 1.**
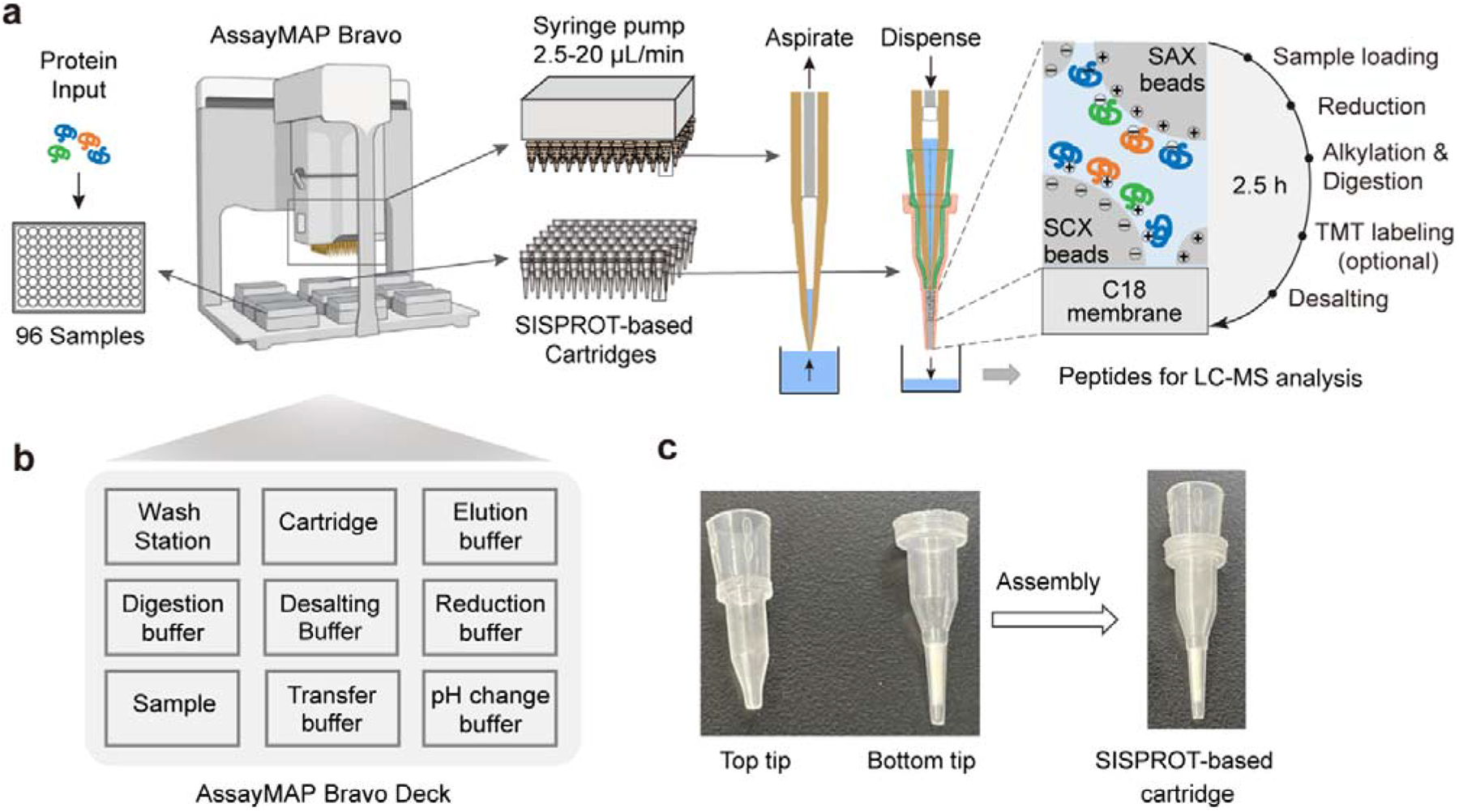
A fully automated and integrated 96-channel sample preparation platform (autoSISPROT) for large-scale proteomics studies. **a** The workflow of autoSISPROT. Protein samples in a 96-well plate are processed by the AssayMAP Bravo workstation, which is equipped with 96-well syringes and SISPROT-based cartridges. All the necessary sample preparation steps, including sample loading, protein reduction, alkylation, digestion, TMT labeling, and desalting, are executed by the programed upward aspiration and downward dispensing of the required buffers through the packed cartridges. The autoSISPROT protocol enables the automatic processing of up to 96 protein samples simultaneously, resulting in peptide solutions within 2.5 hours. **b** The setup of the AssayMAP Bravo deck. **c** The SISPROT-based cartridges are created by assembling top and bottom tips that contain C18 membrane and mixed SCX/SAX beads.

### Performance of autoSISPROT

We benchmarked the sample preparation performance of autoSISPROT platform against manual SISPROT for sample preparation of HEK 293T cell lysates in three technical replicates. We found no significant differences in terms of number of protein groups between autoSISPROT and manual SISPROT, although autoSISPROT identified more peptides than manual SISPROT (**Fig. 2a**). The median coefficient of variation (CV) for autoSISPROT and manual SISPROT was 5.3% and 7.9%, respectively, indicating that autoSISPROT expectedly achieved better quantitative precision (**Fig. 2b**). The comparison of Pearson correlation coefficients also demonstrated better quantitative reproducibility for autoSISPROT (**Fig. 2c**), suggesting that automated operation led to smaller variations. In autoSISPROT samples, 99.7% of cysteine-containing peptides were alkylated, demonstrating a nearly complete reduction and alkylation rection (**Supplementary Fig. 1a**). Furthermore, over 94% of the identified peptides had fewer than two missed cleavage sites, indicating high trypsin digestion efficiency (**Supplementary Fig. 1a**). Additionally, the samples processed by autoSISPROT were subjected to DIA analysis, and the results also showed good reproducibility in identification and quantification (**Supplementary Fig. 1b-d**). Furthermore, we evaluated the TMT labeling performance of autoSISPROT. With autoSISPROT, the TMT-labeled proteins groups and peptides identified from three technical replicates were highly consistent (**Fig. 2d**). Using a cost-effective on-column TMT labeling approach^29^, autoSISPROT achieved high TMT labeling efficiencies for both peptide N-terminus and lysine residues. Approximately 98% of peptide-spectrum matches (PSMs) consistently identified as fully labeled peptides, while the percentage of partially labeled and unlabeled PSMs in three technical replicates was less than 2% (**Fig. 2e**), which aligns with previous studies^30^. Importantly, the fraction of overlabeled PSMs (i.e., off-target TMT labeling on serine, threonine, tyrosine, and histidine) was controlled below 5% (**Fig. 2f**). Collectively, these results demonstrate that autoSISPROT exhibits good sample preparation performance.

**Fig. 2.**
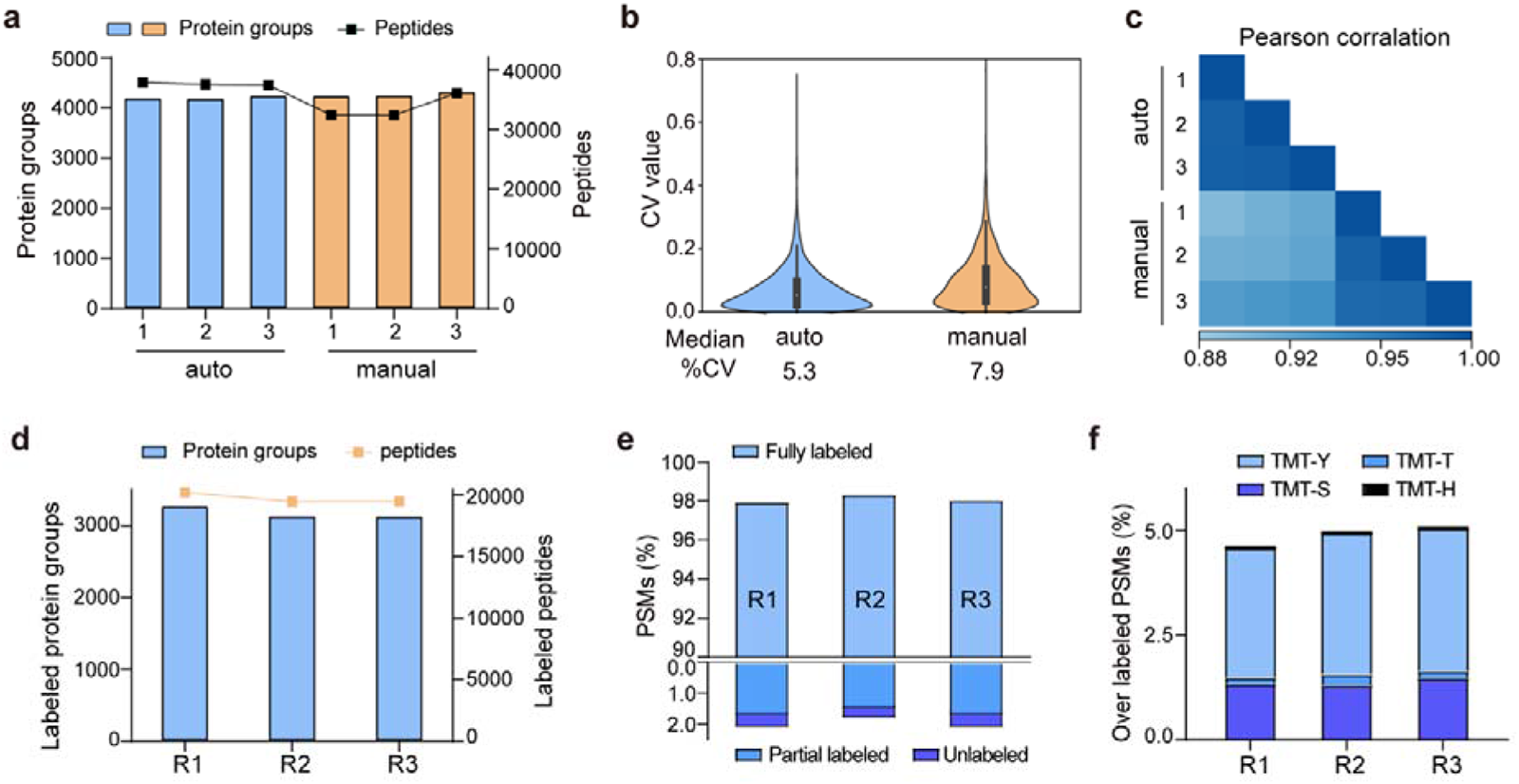
Performance of autoSISPROT. **a** The number of identified protein groups and peptides using autoSISPROT and manual SISPROT under three technical replicates. **b** Violin plots showing the distributions of CVs of protein LFQ intensities between autoSISPROT and manual SISPROT under three technical replicates. **c** Correlation of LFQ intensities of quantified proteins under three technical replicates. **d** Protein groups and peptides identified by autoSISPROT that integrated TMT labeling. **e** TMT labeling efficiency was evaluated using the proportions of fully labeled, partial, and unlabeled PSMs. **f** Overlabeling efficiency was evaluated using the proportions of serine, threonine, tyrosine and histidine labeled PSMs.

### Intra- and inter-batch reproducibility of autoSISPROT

To evaluate the intra- and inter-batch reproducibility of autoSISPROT, we processed 96-well plates with 10 μg of HEK 293T cell lysates per well in three batches on different days, resulting in a total of three 96-well plates and 288 individual samples. For each batch, we randomly selected ten samples for LC-MS/MS analysis (**Fig. 3a**). Consequently, a total of 30 samples were used to evaluate intra- and inter-batch reproducibility of autoSISPROT. Across three batches, an average of 4745 proteins and 80% of zero missed trypsin cleavage sites (digestion performed in room temperature for 1 hour) were identified, with CVs for proteins and zero missed trypsin cleavage sites both less than 3%. (**Fig. 3b**). The intensities distribution of quantified peptides was highly consistent (**Supplementary Fig. 2a**), indicating minimal differences in quantification between intra- and inter-batch analyses. To assess intra-batch reproducibility at the protein level, the CV of proteins quantified with a minimum of three valid values within each batch was calculated. The median CVs for batch 1, batch 2, and batch 3 was 10.8, 12.8, and 12.1%, respectively (**Fig. 3c**), demonstrating highly consistent protein quantification within each batch. The median CVs of intra-batch were comparable to those obtained by autoSP3, which processed 96-wells containing 10 μg of HeLa cell lysates per well^13^. Furthermore, we selected four proteins with different label free quantification (LFQ) intensity range from 10^6^ to 10^10^, and the CVs for these proteins were below 17% across the three batches (**Fig. 3d**). To evaluate inter-batch reproducibility, we calculated Pearson correlation coefficients for proteins with a minimum data completeness of 75% across 30 samples (corresponding to 3403 proteins). Inter-batch comparisons showed highly quantitative reproducibility, with Pearson correlation coefficients exceeding 0.91, indicating no significant differences among the three batches by fully automated sample preparation (**Fig. 3e**). The inset graph in **Fig. 3e** showed a high Pearson correlation coefficient (>0.99) between replicate 5 and replicate 6 of batch 1. As expected, high Pearson correlation coefficients (>0.95) were also achieved for intra-batch comparisons (**Supplementary Fig. 2b-5d**). In summary, these results indicate that autoSISPROT exhibits high intra- and inter-batch reproducibility in sample preparation.

**Fig. 3.**
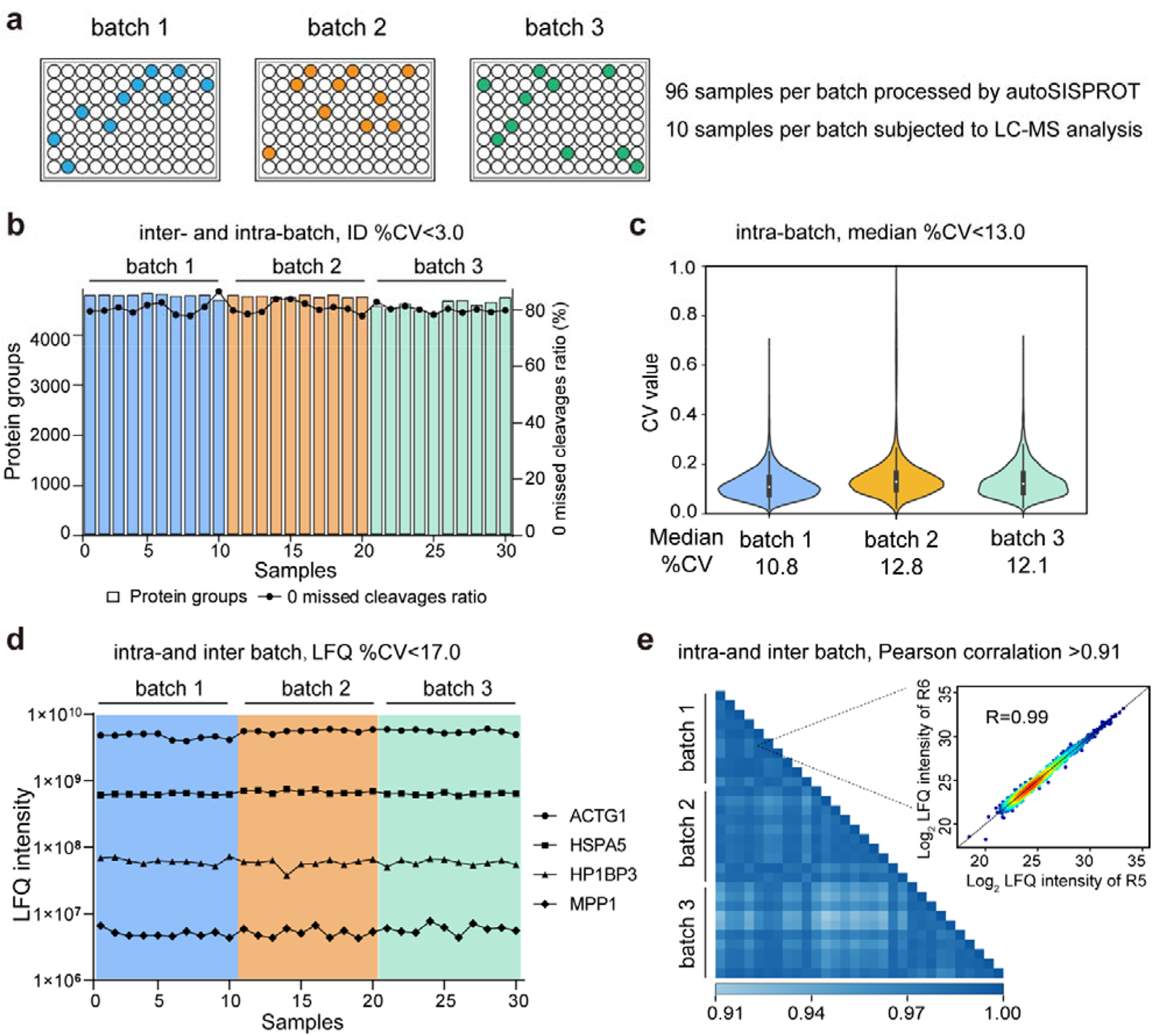
Intra- and inter-batch reproducibility of autoSISPROT. **a** Schematic representation of the experimental design. 96-well plates with 10 μg of HEK 293 cell lysates are processed in three batches at three different days. From each batch, ten randomly selected samples are subjected to LC-MS/MS analysis. **b** Protein groups and percentage of PSMs with zero missed cleavages across three batches. **c** Violin plots depicting CVs distribution of protein LFQ intensities within the same batch. CV values are calculated with a minimum of three valid values within each batch. **d** LFQ intensities of four proteins representing the different dynamic range are plotted across three batches. **e** Pearson correlation coefficient of protein LFQ intensities for inter-batch comparison, and the displayed data are filtered for 75% data completeness^13^. The inset graph showing a correlation between replicate 5 and replicate 6 of batch 1.

### Accurate identification of drug targets by using autoSISPROT

Having established the autoSISPROT protocol, we tested the feasibility of autoSISPROT to identify drug targets via CETSA-MS using MTX as the model drug (**Fig. 4a**). K562 cell lysates were treated with either MTX or a vehicle control, with two independent replicates for each condition, followed by thermal treatment at ten different temperature points. This typical CETSA-MS experimental setting resulted in 40 samples, which would typically take more than one day to obtain TMT labeled peptides ready for analysis by LC-MS by manual operation. Here, the fully automated autoSISPROT significantly reduced the sample preparation time to 3.5 hours. The identified protein groups and peptides were consistent between the two independent replicates per condition (**Fig. 4b**), and high TMT labeling efficiency (> 94%) for both peptide N-terminus and lysine residues were achieved (**Supplementary Fig. 3a**). The boxplot of samples from ten different temperature points exhibited typical sigmoidal trend (**Supplementary Fig. 3b-6e**). Using autoSISPROT, a good correlation (R^2^ = 0.86) of T_m_ assessed with two independent replicates was achieved (**Fig. 4c**), illustrating the high reproducibility of autoSISPROT. As expected, the known target of MTX, dihydrofolate reductase (DHFR), was accurately identified. The melting curves revealed significant changes in DHFR’s thermal stability between the MTX and vehicle treatment conditions (**Fig. 4d**). The shift of melting point (ΔT_m_) of DHFR for two independent replicates were very similar (20.4 °C and 18.4 °C, respectively). Furthermore, the highly consistent melting curves from the two independent replicates per condition indicate the high reproducibility of sample preparation using autoSISPROT. As expected, DHFR ranked first based on the score computed by ProSAP software^31^, which combines the significance of ΔT_m_ and the goodness of fit. The inset graph also showed that DHFR had the most significant changes in ΔT_m_ (**Fig. 4e**). These results demonstrate that autoSISPROT effectively and accurately identifies drug targets, providing an automated platform for drug target identification without excessive manual intervention.

**Fig. 4.**
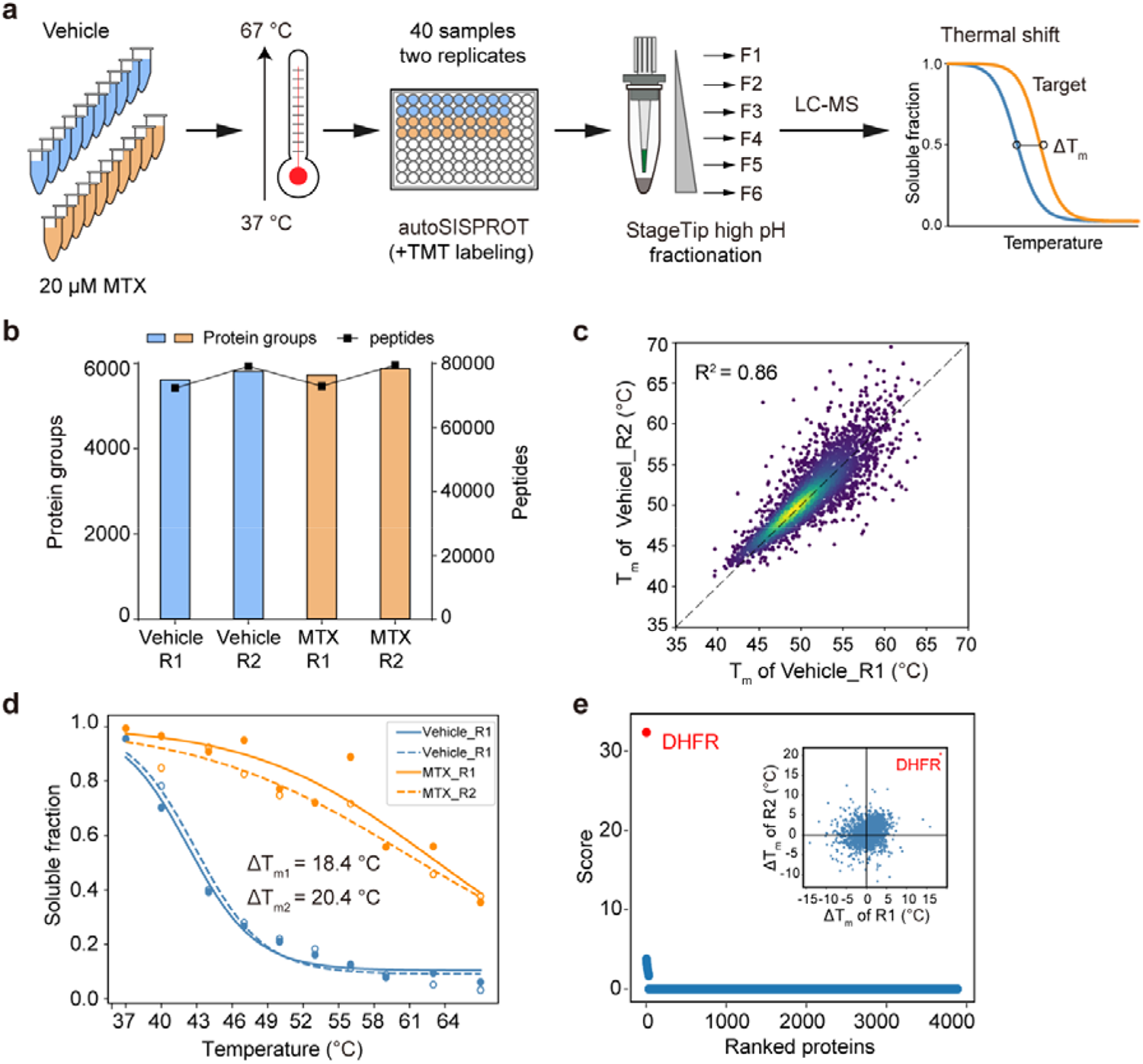
The identification of drug target of MTX by using CETSA and autoSISPROT. **a** Workflow of CETSA and autoSISPROT for drug target identification. **b** Protein and peptide identifications from two independent replicates per condition. **c** Scatter plot showing the correlation of T_m_ between two independent replicates. **d** Melting curves of DHFR in the presence (orange symbols) and absence (blue symbols) of MTX. Data are representative of two independent experiments. **e** Results of drug target identification. Proteins are ranked based on their scores generated by ProSAP software. The inset graph showing scatter plot of ΔT_m_ shifts calculated from the two independent replicates.

### Comparison of TMT and DIA based quantification for CETSA-MS

Current implementations of CETSA-MS typically rely on TMT quantification, and the analysis throughput is limited by the number of TMT channels. In comparison, LFQ with DIA is unlimited in sample numbers and suitable for large-scale drug targets identification. Although DIA has been applied in CETSA-MS analysis, such as mTSA^27^, which has been published at the time of this writing, no studies have systematically evaluated the performance of TMT and DIA methods for CETSA-MS analysis. Here, we conducted a comparison between the TMT and DIA methods, referred to as tmtCETSA and diaCETSA, respectively. We utilized staurosporine, a pan-kinase inhibitor, as the model drug since its targets have been extensively studied on a proteome scale^23, 26, 27^. K562 cell lysates were treated with either staurosporine or a vehicle control and subjected to thermal treatment at 52°C. Afterwards, equal aliquots of the soluble lysates were individually processed using the diaCETSA and tmtCETSA methods (**Supplementary Fig. 4a**). To ensure a fair comparison, the number of samples, MS instrument, and MS acquisition time were kept the same for both methods. In comparison to classical CETSA-MS, diaCETSA exhibited a 4-fold improvement in throughput by reducing the number of samples from 40 to 10, while also saving time and costs (particularly for TMT labeling and peptide fractionation) compared to tmtCETSA. Within our study, tmtCETSA resulted in 20-30% more proteins compared to diaCETSA, primarily due to the benefit of fractionation (**Supplementary Fig. 4b**), which was consistent with previous work^32^. Both methods achieved a high Pearson correlation coefficient (> 0.98) and good quantitative precision (median CVs < 10%) at the protein level (**Supplementary Fig. 4c-7d**), demonstrating good quantitative reproducibility for both methods. Furthermore, using tmtCETSA and diaCETSA, we identified 120 and 106 proteins, respectively, as staurosporine targets that met the significance criteria (adjusted p-value < 0.05). The diaCETSA method identified slightly fewer kinase targets (53 kinases) compared to tmtCETSA (67 kinases) (**Supplementary Fig. 4b**), which could be attributed to the lower proteomic depth of diaCETSA. Subsequently, we benchmarked our dataset against classical TPP. Overall, 24 kinase targets were found to be shared among these three methods (**Supplementary Fig. 4f**).

Interestingly, we found that certain non-kinase proteins were identified as targets by both methods. String analysis revealed that these non-kinase proteins had interactions with kinases, suggesting that indirect interactions with staurosporine could be identified by thermal stability analysis (**Supplementary Fig. 4g-7h**, left panel). Enrichment analysis of Gene Ontology (GO) molecular functions of the significant proteins revealed their involvement in various kinase activities, suggesting that these proteins may play functional roles in different kinase-related processes (**Supplementary Fig. 4g-7h**, middle panel). Furthermore, we mapped the kinase targets identified by both methods onto the kinome tree (**Supplementary Fig. 4g-7h**, right panel). Notably, we observed that several kinases, such as PRKCB, PRKCD, PRKCI, and SLK, exhibited destabilization upon staurosporine treatment, as identified by both tmtCETSA and diaCETSA.

We further optimized the diaCETSA method to enhance the depth of proteome analysis. Given the significant impact of the spectral library’s quality on DIA data analysis, we initially compared two spectral libraries constructed from K562 cell lysates using different sample preparation approaches. Spectral library 1 was generated using a conventional proteomics workflow, involving protein precipitation and in-solution digestion^33^. On the other hand, samples for constructing spectral library 2 underwent heat treatment at 52°C and were processed using autoSISPROT. Despite Library 1 having a larger capacity (protein number: 9589 versus 6000), more proteins were identified when searching with Library 2 (**Supplementary Fig. 5**). This demonstrated that the project-specific library does not need to be as extensive as possible but should be built in a manner consistent with the analyzed DIA samples. Additionally, we compared the quantitative performance of DIA mode on two different mass spectrometers, i.e., Orbitrap Exploris 480 and timsTOF Pro, and found that Orbitrap Exploris 480 exhibited slightly better protein quantification and identified more drug targets (**Supplementary Fig. 6**). Overall, we employed these optimal conditions for the subsequent diaCETSA analysis.

### Combining autoSISPROT and diaCETSA for high-throughput identification of drug targets and off-targets

To evaluate whether autoSISPROT could achieve high-throughput drug target identification, we applied autoSISPROT coupled with diaCETSA to identify targets of 24 drugs including 20 kinase inhibitors (KIs) (**Fig. 5a**). In diaCETSA, different drug treatment conditions can share a common vehicle control, which further improves the analysis throughput. These 24 drugs were classified into two groups based on whether they had been previously studied using CETSA-MS methods and were processed on different days. Compared to the classical CETSA-MS, our method provided a more than 10-fold improvement in throughput (960 CETSA-MS samples versus 87 diaCETSA samples for 24 drugs). The number of identified protein groups and enzymatic cleavage efficiency were highly consistent across all sets of samples (**Supplementary Fig. 7a-d**). In addition, median CVs of LFQ intensities were < 25% for all 24 drugs (**Supplementary Fig. 7e, f**), demonstrating good processing precision spanning the entire sample preparation procedure from thermal treatment to MS data acquisition. Among the 20 kinase inhibitors, we successfully identified the reported targets for 12 kinase inhibitors (**Fig. 5a**). The annotated kinome tree shows that these identified kinase targets mostly belong to protein kinase families of GCMC, CAMK, and AGC (**Fig. 5b**). Furthermore, our method can effectively identify the known targets and yielded complementary results with previously published methods for palbociclib and dinaciclib (**Fig. 5c**).

**Fig. 5.**
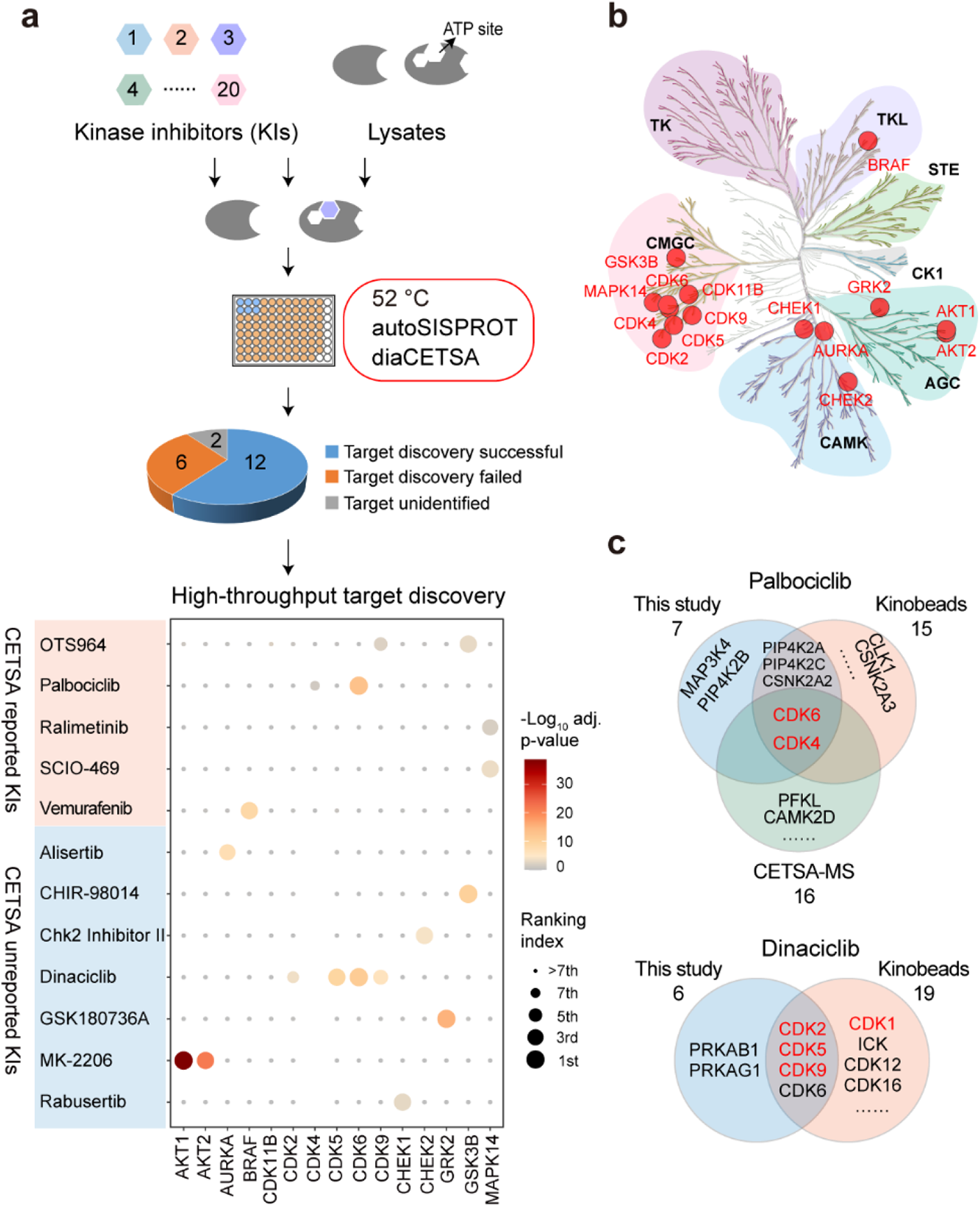
High-throughput drug target identification for kinase inhibitors by combing autoSISPROT and diaCETSA. **a** Workflow for high-throughput identification of targets of kinase inhibitors. Dot plots visualization of targets identification of OTS964, palbociclib, ralimetinib, SCIO-469, vemurafenib, alisertib, CHIR-98014, Chk2 Inhibitor II, dinaciclib, GSK180736A, MK-2206, and rabusertib. **b** Kinome tree displaying all identified kinase targets**. c** Venn diagram displaying the common and complementary targets of palbociclib and dinaciclib identified by kinobeads, classical CETSA-MS, and our method. The proteins marked in red are the known targets.

We successfully identified the reported targets for all of the five CETSA-studied KIs, namely OTS964, palbociclib, ralimetinib, SCIO-469, and vemurafenib and seven CETSA-unstudied KIs (**Fig. 5a**). Notably, all of the identified kinase targets ranked within the top seven hits, except for OTS964, indicating the high accuracy of our method. Moreover, we accurately identified the targets of four CETSA-studied non-kinase inhibitors: methotrexate, olaparib, panobinostat, and raltitrexed. This further demonstrates the robustness of the autoSISPROT and diaCETSA methods. Regarding the CETSA-unstudied KIs, however, target identification could fail due to several reasons, such as the absence of thermal stabilization effect or the low abundance of target proteins in K562 cells. Our results show that the known targets of six KIs (bafetinib, bosutinib, BS-181 tideglusib, SGC-GAK-1, and roscovitine) did not exhibit significant differences between the drug and vehicle treatment conditions. To determine whether this failure was due to the lack of a thermal stabilization effect or the limitations of the single temperature point strategy, we conducted classical CETSA-MS analysis using ten temperature points to identify the drug targets of SGC-GAK-1. However, even with the increased temperature points, the known target of SGC-GAK-1 (cyclin-G-associated kinase, GAK) did not show significant changes between the drug and vehicle treatment conditions. As for the other two KIs (KN-62 and saracatinib), the abundance of their known targets (CaMK2 and SRC) in K562 cells was too low to be determined.

Based on the principle of CETSA, our method also enabled the high-throughput identification of potential drug off-targets. For instance, we also identified the reported off-targets of dinaciclib (CDK6), palbociclib (PIP4K2A, PIP4K2C, and CSNK2A2)^34^ and vemurafenib (FECH and MAP2K4), along with their known targets. In addition, our results reveal several new potential off-targets for kinase inhibitors (e.g., OTS964, vemurafenib, alisertib, dinaciclib, etc.). Overall, these results demonstrate that the combination of autoSISPROT and diaCETSA enables the identification of drug targets and off-targets in a fully automated manner and is well-suited for high-throughput drug target identification.

## Discussion

The SISPROT technology allows for the seamless integration of multiple steps in protein sample preparation, including preconcentration, reduction, alkylation, digestion, desalting, and high-pH RP-based peptide fractionation, all within a single spintip device. Its foundation on on-bead digestion, known for its high efficiency, makes it easily adaptable to robotic automation. Most of the currently reported automated and high-throughput sample preparation methods rely on in-solution digestion strategies using robotic liquid handling workstations like Bravo, making it challenging to achieve fully automated and integrated sample preparation. In contrast, autoSISPROT enables fully automated and integrated 96-well sample preparation by performing on-bead digestion in the AssayMAP Bravo system. Through the seamless integration of all sample preparation steps into self-designed cartridges, the end-to-end autoSISPROT method is rapid, completing the processing of 96 samples from protein input to peptide elution in just 2.5 hours. No manual operations are required after clicking the start button when performing autoSISPROT, thereby significantly reducing multi-step manual operations, hands-on time, and the variability in protein quantification. By integrating the on-column TMT labeling step into the automated protocol, autoSISPROT only takes 3.5 hours to complete the digestion and TMT labeling of 96 samples, providing a streamlined solution for automated and reproducible quantitative proteomics. We demonstrated the excellent performance of autoSISPROT in sample preparation, and importantly, we also showcased its good intra- and inter-batch reproducibility. This was evidenced by CVs for protein identification and quantification below 3.0% and 17%, respectively, as well as Pearson correlation coefficients of more than 0.95 and 0.91 for intra- and inter-batch analyses, respectively.

Notably, the cartridge used in autoSISPROT was specifically self-designed, taking reference from a standard 200 μL tip. The cartridge had several key requirements: (1) The inner part of the top tip outlet had a similar geometry to syringes, allowing the cartridges to be picked up by syringes while maintaining air tightness and avoiding leakage. (2) The overfill position of the bottom tips had a similar geometry to the housing of the top tip outlet, enabling them to be assembled in an overfill manner to ensure air tightness. (3) The cartridges needed to be sufficiently stable to withstand the back pressure exerted during packing with SISPROT materials and organic solvents like ACN. (4) The cartridges should be cost-effective and disposable. Industry-level moldmaking technology and polypropylene (PP) material were employed for fabricating the cartridges, allowing for mass production of thousands of cartridges at a cost of less than 0.3 US dollars per cartridge.

CETSA-MS has been widely used for identifying targets and off-targets of various drugs. Currently, CETSA-MS and improved throughput methods such as PISA, iTSA, and mTSA rely on laborious manual sample preparation, making high-throughput drug target identification challenging. Encouragingly, the automated sample preparation platforms developed in this study provide multifunctional options for CETSA-based drug target identification. By combining autoSISPROT and CETSA-MS technology, we selectively identified the known target of MTX in an automated manner. Additionally, the combination of autoSISPROT and diaCETSA allowed for the identification of kinase targets for 20 drugs in a fully automated and high-throughput manner. Compared to manual CETSA-MS, our automated platform significantly reduced manual operation and hands-on time while improving analysis throughput. By incorporating autoSISPROT and diaCETSA, up to 127 drugs can be analyzed in a single automated operation using 384-well plates, greatly facilitating proteomics-based drug discovery. Collectively, as large-cohort proteomic analysis continues to advance, autoSISPROT will provide a multifunctional and end-to-end solution for automated, robust, and reproducible sample preparation without manual intervention.

## Methods

### Cell culture and protein extraction

The cancer cell lines HEK 293T and K562 were purchased from American Type Culture Collection. HEK 293T cells were cultured in Dulbecco’s modified Eagle’s medium (Corning), supplemented with 10% fetal bovine serum (Gibco), 100 U/mL penicillin (Invitrogen), and 100 μg/mL streptomycin (Invitrogen) in a humidified incubator with 5% CO_2_ at 37 °C. HEK 293T Cells were harvested at ∼80% confluence by washing three times with phosphate buffered saline (PBS) buffer and lysed in the lysis buffer containing 20 mM HEPES, pH 7.4, 150 mM NaCl, 600 mM guanidine HCl, 1% n-dodecyl β-D-maltoside (DDM), and protease inhibitors (1 mM PMSF, 1 μg/mL leupeptin, 1 μg/mL pepstatin, and 1 μg/mL aprotinin). The obtained cell lysates were sonicated and centrifuged at 18000 × g for ∼30 min at 4 °C. The protein concentration was measured using the BCA assay (Thermo Fisher Scientific, Germany), and the final protein concentration was adjusted to 5 mg/mL.

K562 cells were cultured in RPMI 1640 medium (Gibco). Cells were harvested by washing three times with PBS buffer. For CETSA experiments, the K562 cell pellet was resuspended in a lysis buffer containing a final concentration of 50 mM HEPES (pH 7.4), 5 mM β-glycerophosphate, 0.1 mM activated Na_3_VO_4_, 10 mM MgCl_2_, 1 mM tris(β-chloroethyl) phosphate (TCEP, Sigma), and EDTA-free protease inhibitor (Roche) and lysed by three rounds of flash-freeze-thaw cycles (alternating exposure of the samples to liquid nitrogen and 37 °C in a water bath). Mechanical shearing was carried out by passing the thawed suspension through a syringe with a narrow needle several times. The resulting cell lysates were pelleted by centrifugation at 18,000 × g for approximately 30 min at 4 °C, and the supernatant was collected. The final protein concentration was adjusted to 5 mg/mL as determined by the BCA assay. For building the DDA spectral library, the K562 cell pellet was lysed in a lysis buffer containing 8 M urea, 50 mM ammonium bicarbonate (ABC), and a protease inhibitor mixture. The subsequent steps for protein extraction were the same as those for the HEK 293T cell lysate.

### Thermal treatment of K562 cell lysates

The procedure for the ten temperature points-based CETSA-MS experiments followed a reported protocol^23^. Briefly, K562 cell lysates were divided into ten identical aliquots (100 μg per aliquot, 20 μL) and treated with either 20 μM drug in 1% DMSO or with DMSO alone. With two replicates per condition, a total of 40 treated samples were incubated at room temperature for 5 min before heat treatment. Subsequently, all samples were transferred to a PCR machine and heated for 3 min at ten different temperature points (37, 40, 44, 47, 50, 53, 56, 59, 64, and 67 °C), followed by a 5-minute incubation at 4 °C. After heat treatment, the protein aggregates were removed by centrifugation at 18000 × g for approximately 30 min at 4 °C, and the supernatant was collected.

For ten temperature points based CETSA-MS experiments, thermal treatment was performed similarly to the procedure as reported by previous work^23^. Briefly, K562 cell lysates were split into ten identical aliquots (100 μg per aliquot, 20 μL) and treated with either 20 μM drugs in 1% DMSO, or with DMSO alone. With two replicates per condition, 40 treated samples were incubated at room temperature for 5 min prior to heat treatment. After incubation, all samples were transferred to a PCR machine and heated for 3 min to ten different temperature points (37, 40, 44, 47, 50, 53, 56, 59, 64, and 67 °C), followed by a 5 min incubation at 4 °C. After heat treatment, the protein aggregates were removed by centrifugation at 18000 × g for 30 min at 4 °C, and the supernatant was collected.

For the single temperature point-based CETSA experiments, K562 cell lysates were divided into two identical aliquots (100 μg per aliquot, 20 μL), treated with 20 μM drug in 1% DMSO, or with DMSO alone. In the tmtCETSA versus diaCETSA comparison experiment, five replicates per condition were chosen. For high-throughput diaCETSA experiments, six replicates were chosen for the vehicle condition and three replicates for the drug condition, with multiple drug conditions sharing the common vehicle conditions. The treated samples were incubated at room temperature for 5 min and then transferred to a PCR machine for 3 min of heating at 52 °C, followed by a 5-minute incubation at 4 °C. The remaining steps for thermal treatment were the same as described above.

### Fabrication and screening of the SISPROT-based cartridges

The preparation of SISPROT-based cartridges strictly followed ten steps. In brief, the SISPROT-based cartridges were constructed by assembling the top and bottom tips. The bottom tips were packed sequentially with three plugs of C18 membrane (3 M Empore) and mixed-mode ion exchange beads (SCX:SAX = 1:1, Applied Biosystems). The quantity of C18 membrane and mixed-mode ion exchange beads was adjusted based on the protein amount.

### Sample preparation for building DDA spectral library

For the construction of DDA spectral library 1, K562 cell lysates underwent in-solution trypsin digestion following a previous study^33^. Prior to the digestion reaction, proteins were purified using methanol-chloroform precipitation^35^. The resulting peptides were desalted using a Sep-Pak Vac 1cc tC18 cartridge (Waters), dried using SpeedVac, and stored at -20 °C until LC-MS/MS analysis. For building DDA spectral library 2, sample preparation was processed by autoSISPROT method, as described below.

### Sample preparation by using autoSISPROT

The autoSISPROT method was carried out using an AssayMAP Bravo equipped with 96-well ultra-low dead volume syringes and the homemade disposable SISPROT-based cartridges.

The autoSISPROT workflow involved eleven key steps, which were (1) activation of SISPROT-based cartridges with activation buffer (Deck 3), (2) equilibrium of SISPROT-based cartridges with equilibrium buffer (Deck 5), (3) loading acidified cell lysates (pH 2.0-3.0, Deck 7) to SISPROT-based cartridges, (4) washing SISPROT-based cartridges with activation buffer, (5) washing SISPROT-based cartridges with equilibrium buffer, (6) reduction of disulfide bonds with reduction buffer (Deck 6) for 30 min at room temperature, (7) washing with pH change buffer (Deck 9) to adjust pH, (8) loading alkylation and digestion buffer (Deck 4) for 60 min at room temperature (in darkness), (9) transferring digested peptides from the mixed-mode ion exchange beads onto the C18 membrane with transfer buffer (Deck 8), (10) desalting peptides with desalting buffer, and (11) eluting peptides with elution buffer. The eluted peptides were collected into 96-well plates. Finally, the samples were dried using SpeedVac and stored at -20 °C before LC-MS/MS analysis. During the autoSISPROT operation, no manual operations were required once all buffers were transferred to the corresponding 96-well plates.

For TMT-based quantitative proteomics analysis, the buffer in Deck 9 is replaced with HEPES buffer. Then two additional steps are implemented: pH adjustment using HEPESbuffer and TMT labeling using a labeling buffer containing 0.4 μg/μL of TMT reagents in HEPES buffer. Moreover, a pause step is necessary to replace the sample plate with the TMT reagent plate before TMT labeling.

### Sample preparation by using manual SISPROT

We conducted manual SISPROT using the same SISPROT-based cartridges, buffers, and HEK 293T cell lysates as in autoSISPROT. However, in the manual method, all sample preparation steps were carried out using centrifugation, following the procedures described in previous studies^10, 11^.

In the tmtCETSA versus diaCETSA comparison experiment, we employed the reported mixed-mode SISPROT for sample preparation with slight modifications^11^. After digestion, ten samples from the tmtCETSA experiment were labeled with a TMT10-plex kit (Thermo Fisher Scientific, Germany) using the TMT-based FISAP method^29^. Subsequently, the eluants from the tmtCETSA experiments were combined into a single sample, dried using SpeedVac, and stored at -20 °C before fractionation. On the other hand, ten samples from the diaCETSA experiment were directly eluted without labeling, dried using SpeedVac and stored at -20 °C before LC-MS/MS analysis.

### High-pH reversed phase fractionation

To generate a comprehensive DDA spectral library of K562 cells, peptide fractionation was conducted using a 1260 Infinity II HPLC system (Agilent Technologies) with a XBridge peptide BEH C18 column (2.1 mm i.d. × 150 mm) at a flow rate of 200 μL/min. Buffer A consisted of 2% ACN in 10 mM ABC (pH 8.0), while buffer B comprised 90% ACN in 10 mM ABC (pH 8.0). Peptides were separated using a 70-minute segmented gradient as follows: 1%-9% buffer B in 1 minute, 9%-35% buffer B in 50 min, 35%-70% buffer B in 4 min, followed by a 15-minute wash with 70% buffer B. Fractions were collected every 30 seconds and combined into 24 fractions.

In the tmtCETSA versus diaCETSA comparison experiment, the TMT-labeled peptide mixture was separated using a 40-minute segmented gradient as follows: 1%-13% buffer B in 1 minute, 13%-52% buffer B in 29 min, 52%-90% buffer B in 4 min, followed by a 6-minute wash with 90% buffer B. Fractions were collected every 30 seconds and combined into 10 fractions. The obtained fractions were dried using SpeedVac and stored at -20 °C before LC-MS/MS analysis.

For the ten temperature points based CETSA-MS experiments, the TMT-labeled peptide mixture underwent fractionation using C18 StageTip-based high-pH reversed-phase fractionation^29^. The peptide mixture was fractionated using 10 μL portions of 18 different elution buffers (3%, 5%, 7%, 9%, 11%, 13%, 15%, 17%, 19%, 21%, 23%, 24%, 26%, 28%, 30%, 35%, 40%, and 80% ACN) in 5 mM ABC at pH 10.0. Subsequently, 18 fractions were combined into six fractions, dried using SpeedVac, and stored at -20 °C.

### LC-MS/MS analysis

Nano-flow LC-MS/MS was performed on an Orbitrap Exploris 480 equipped with an UltiMate 3000 (Thermo Fisher Scientific, Germany), or on a timsTOF Pro equipped with a nanoElute (Bruker Daltonics, Germany). Buffer A was 0.1% FA in water for both Dionex UltiMate 3000 and nanoElute, and buffer B was 0.1% FA in 80% ACN for Dionex UltiMate 3000 and 0.1% FA in 100% ACN for nanoElute. Chromatographic separation was performed via the homemade 100 μm i.d. × 20 cm analytical column packed with 1.9 μm/120 Å C18 beads (Dr. Maisch GmbH, Germany) at a flow rate of 500 nL/min (Dionex UltiMate 3000) or 300 nL/min (nanoElute). When using the Dionex UltiMate 3000 system to separate unlabeled peptides, an 85-minute segmented gradient was applied as follows: 4%-8% buffer B in 2 min, 8%-28% buffer B in 53 min, 28%-36% buffer B in 10 min, 36%-100% buffer B in 9 min, followed by an 11-minute equilibration with 1% buffer B. For separating TMT-labeled HEK 293T cell peptides, a 75-minute segmented gradient was used: 4%-9% buffer B in 2 min, 9%-35% buffer B in 51 min, 35%-50% buffer B in 10 min, 50%-100% buffer B in 3 min, followed by a 3-minute wash with 100% buffer B and a 6-minute equilibration with 1% buffer B. For the analysis of unlabeled K562 cell peptides, peptide samples were spiked with iRT peptides from Biognosys (Switzerland) for retention time calibration. A 135-minute segmented gradient was used: 4%-8% buffer B in 2 min, 8%-28% buffer B in 105 min, 28%-40% buffer B in 15 min, 40%-99% buffer B in 1 minute, followed by a 5-minute wash with 99% buffer B and a 7-minute equilibration with 1% buffer B. For analyzing TMT-labeled K562 cell peptides, a 135-minute segmented gradient was employed: 4%-9% buffer B in 2 min, 9%-35% buffer B in 103 min, 35%-50% buffer B in 15 min, 50%-100% buffer B in 3 min, followed by a 5-minute wash with 100% buffer B and a 7-minute equilibration with 1% buffer B. When using the nanoElute system to separate unlabeled K562 cell peptides, an 80-minute segmented gradient was applied: 2%-24% buffer B in 50 min, 24%-36% buffer B in 10 min, 36%-80% buffer B in 10 min, followed by a 10-minute wash with 100% buffer B.

Orbitrap Exploris 480 instrument was operated in positive ion mode with the following settings: an electrospray voltage of 2.0 kV, a funnel RF lens value of 40, and an ion transfer tube temperature of 320 °C. MS1 scans were performed in the orbitrap analyzer, covering an m/z range of 350 to 1200, with a resolution of 60000. The automatic gain control (AGC) target value was set to 3E6, and the maximum injection time (MIT) was in auto mode. The MS/MS spectra were acquired using DDA mode, with one MS scan followed by 40 MS/MS scans. Precursors were isolated by the quadrupole using a 1.4 Da window, followed by higher-energy collisional dissociation (HCD) fragmentation using a normalized collision energy (NCE) of 30%. Fragment ions were scanned at a resolution of 7500. AGC target and MIT for MS2 scans were set to standard mode and 15 ms, respectively. The scanned peptides were dynamically excluded for 30 s. Monoisotopic precursor selection was enabled, and peptide charge states from +2 to +6 were selected for fragmentation.

For TMT-labeled peptides, MS1 scans were performed in the orbitrap analyzer, covering an m/z range of 350 to 1400, with an MIT of 45 ms for MS1 scans. The instrument was set to run in top speed mode with 2 s cycles for the survey and MS/MS scans. HCD fragmentation was performed with a NCE of 36%. Fragment ions were scanned at a resolution of 30000, with an AGC target of 1E5 and an MIT of 54 ms. TurboTMT was operated in TMT reagent mode, with a precursor fit threshold of 65% and a fit window of 0.7 Da.

For DIA mode, each MS1 scan was followed by 60 variable DIA windows with 1.0 Da overlap, and the remaining parameter settings were the same as described above. In the tmtCETSA versus diaCETSA comparison samples, the FAIMS Pro device from Thermo Fisher Scientific (Germany) was used. The FAIMS device parameters included an inner electrode temperature of 100 °C, an outer electrode temperature of 100 °C, a carrier gas flow rate of 0 L/min, an asymmetric waveform with a dispersion voltage of -5000 V, and an entrance plate voltage of 250 V. The selected CV (-45 and -65 V) was applied throughout the LC-MS/MS run for static CV conditions.

Bruker timsTOF Pro was operated in positive ion mode using a captive nano-electrospray source at 1500V. The MS operated in DDA mode for ion mobility-enhanced spectral library generation. The accumulation and ramp time for mass spectra were both set to 100 ms, and the recorded mass spectra ranged from m/z 300 to 1500. Ion mobility was scanned from 0.75 to 1.40 Vs/cm^2^. The overall acquisition cycle consisted of one full TIMS-MS scan and 10 parallel accumulation-serial fragmentation (PASEF) MS/MS scans. During PASEF MS/MS scanning, the collision energy was linearly ramped as a function of mobility from 59 eV at 1/K_0_=1.40 Vs/cm^2^ to 20 eV at 1/K_0_=0.75 Vs/cm^2^. In DIA mode, 32 × 25 Da isolation windows were defined from m/z 400 to 1200. To adapt the MS1 cycle time in diaPASEF, the repetitions were set to 2 in the 16-scan diaPASEF scheme.

### Data analysis

The raw data were searched with MaxQuant (version 1.6.2.3) against a human protein database (release 2020_03, 74811 entries). Unless otherwise noted, the same database was used and the default parameters were employed. Trypsin was set as the enzyme with up to two missed cleavages. Carbamidomethylation (+57.021 Da) of cysteine was set as a fixed modification, and protein N-terminal acetylation (+42.011 Da) and oxidation of methionine residues (+15.995 Da) were considered as variable modifications. The false discovery rate (FDR) was set to 1% at the site, peptide-spectrum match (PSM), and protein levels. For TMT-labeled samples, Proteome Discoverer™ software (version 2.4.1.15) was used for the search. Precursor mass tolerance was set to 10 ppm, and fragment ions were set to 0.02 Da. TMT tags on lysine residues and peptide N-terminus (+229.163 Da) were defined as static modifications, while the other fixed and variable modification remained consistent with those mentioned above. PSMs were validated using the Percolator algorithm, peptides were validated using the Peptide Validator algorithm, and proteins were validated using the Protein FDR Validator algorithm. Proteins were quantified by summing reporter ion counts across all matching PSMs. DIA raw data were searched with Spectronaut (version 15.0) with default parameters.

The output results from MaxQuant were used to generate density plots, heat maps, boxplots, and dot plots using R (version 3.4.0). The output results from Proteome Discoverer or Spectronaut were used to generate violin plots, melting curves, scatter plots of T_m_ and ΔT_m_ shifts using Python (version 3.9). All the volcano plots were created using the ProSAP^31^ software and the p-value was calculated to assess the statistical significance of T_m_ after a Benjamini-Hochberg correction. Molecular function annotation was based on the GO knowledgebase (http://geneontology.org/). Protein-protein relationships were analyzed using STRING (version 11.5; https://string-db.org/). Kinome trees were built using the kinmapbeta tool (http://www.kinhub.org/kinmap/)^36^.

## Acknowledgements

This work is supported by grants from the China State Key Basic Research Program Grants (2021YFA1301601, 2021YFA1302603, 2021YFA1301602, 2020YFE0202200, and 2022YFC3401104), the National Natural Science Foundation of China (22125403, 22074060, 22150610470, and 92253304), the Shenzhen Innovation of Science and Technology Commission (JCYJ20200109140814408, JCYJ20210324120210029 and JCYJ20200109141212325), and Guangdong province (2019B151502050).

## Author contributions

Q.W., X.S., J.Z., R.T. and C.T. designed the experiments. Q.W. and X.S. performed the sample preparation and proteomics experiments. J.Z. and C.F. conducted biophysical, cell biology, and molecular biology experiments. X.C. and X.L. provided assistance in the preparation of SISPROT-based cartridges. Q.W., H.J., Y.L., A.H., and B.L. analyzed the data. R.T. conceived and supervised the project. Q.W., J.Z., C.T. and R.T. wrote the paper.

## Competing Interests

The authors declare no competing interests.

## Notes

### Competing Interest Statement

R.T. is a founder for BayOmics, Inc. The other authors declare no competing interests.

