## Supplementary information for "Fully automated and integrated proteomics sample preparation platform for high-throughput drug target identification"

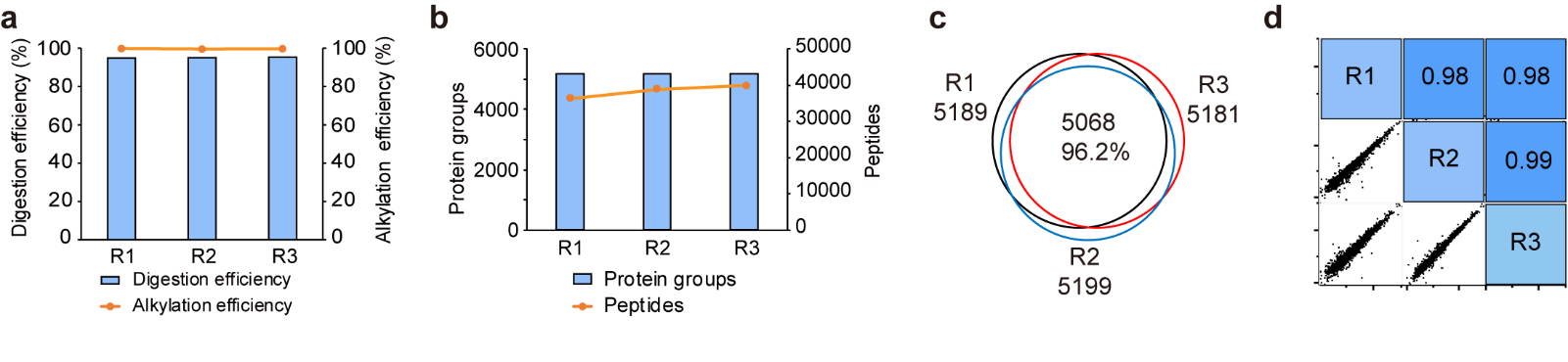

**Supplementary Fig. 1 Performance of the proteome profiling by using autoSISPROT**. **a** Alkylation and digestion efficiencies of autoSISPROT for processing 10 μg of HEK 293T cell lysates under three technical replicates. **b** The number of protein groups and peptides identified with DIA. **c** The number of common protein groups identified by autoSISPROT identified with DIA under three technical replicates. **d** Correlation of LFQ intensities of proteins quantified with DIA under three technical replicates.

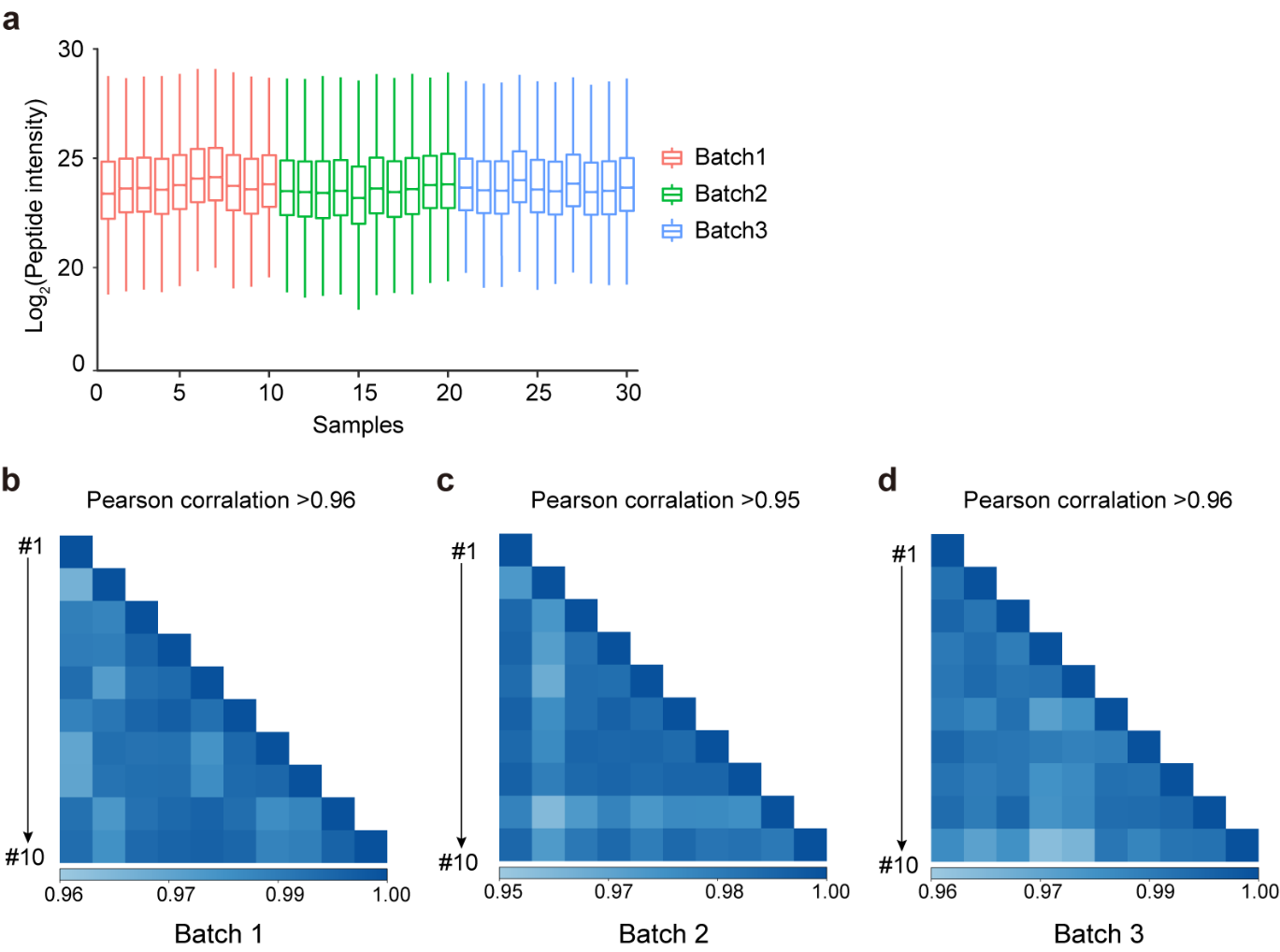

**Supplementary Fig. 2** **Evaluation of intra- and inter-batch quantified reproducibility of autoSISPROT**. **a** Boxplots of log2-transformed peptide intensities across three batches samples. The color coding highlights the samples batch of origin. **b-d** Pearson correlation coefficient of protein LFQ intensities for (**b**) batch 1, (**c**) batch 2, and (**d**) batch 3, respectively.

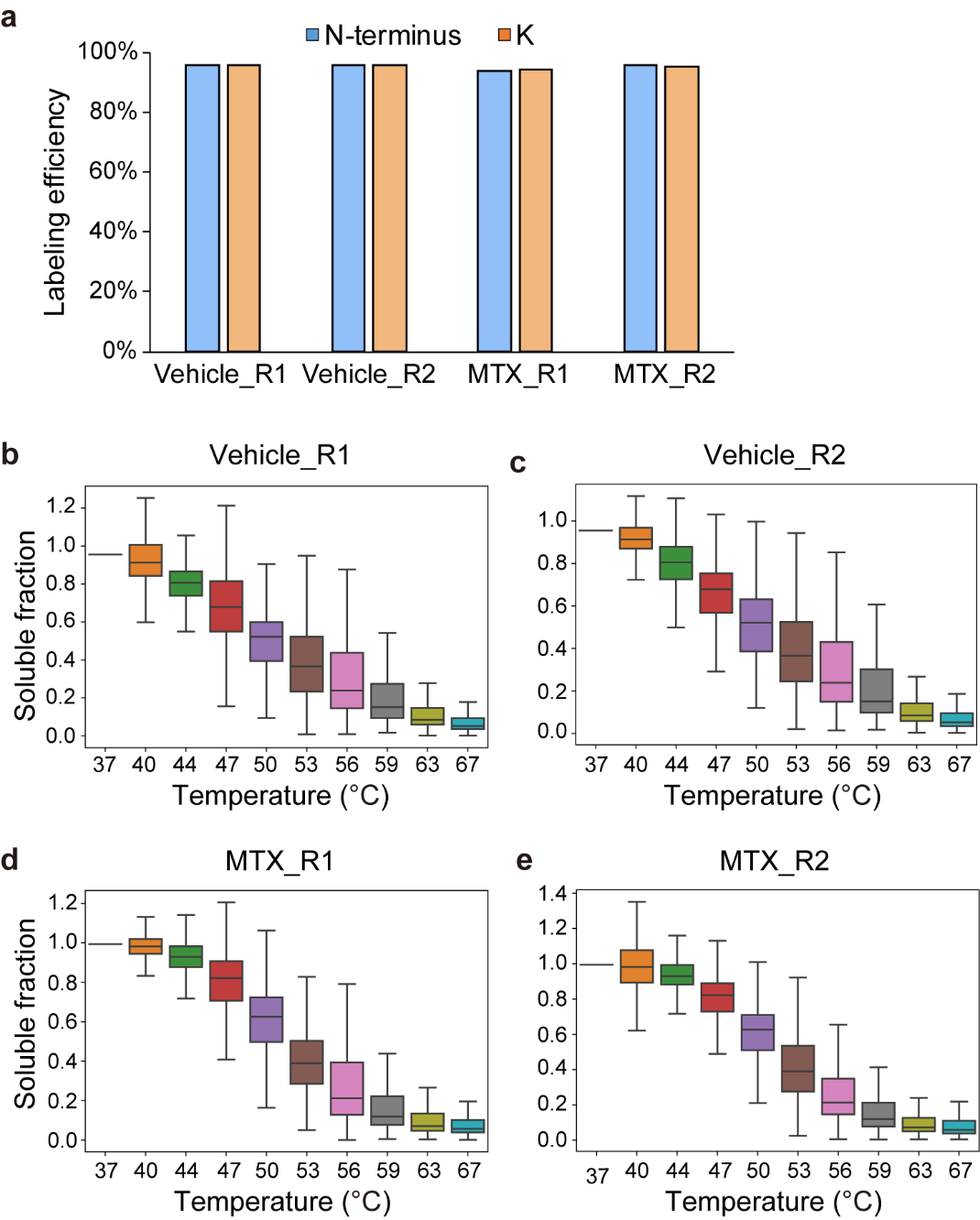

**Supplementary Fig. 3 Performance of autoSISPROT for processing the samples from TMT-based CETSA-MS**. **a** TMT labeling efficiency of peptide N-terminus and lysine residues for CETSA samples. **b-e** Boxplot of soluble fraction from indicated temperatures for (**b**) Vehicle_R1, (**c**) Vehicle_R2, (**d**) MTX_R1, and (**e**) MTX_R2, respectively.

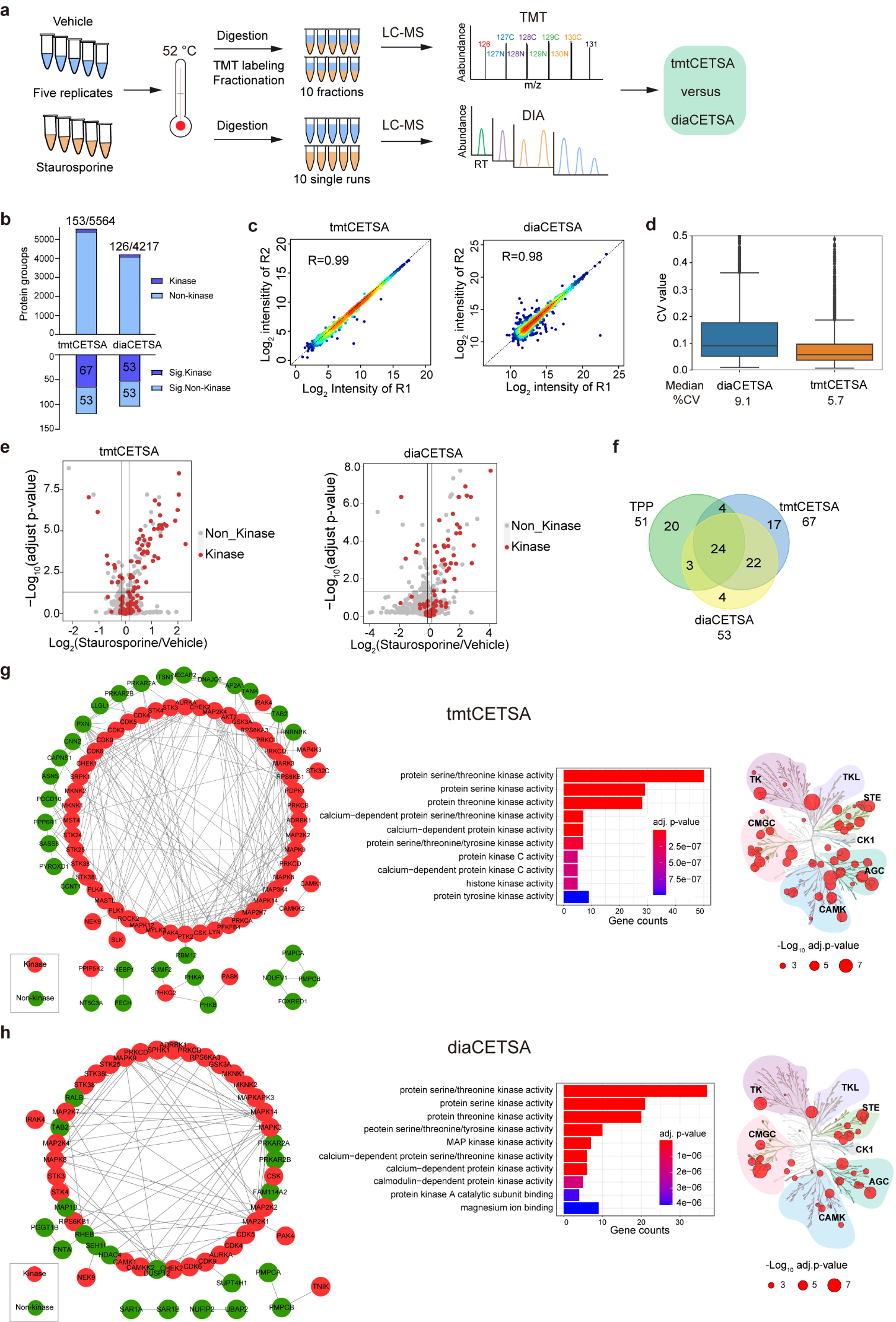

**Supplementary Fig. 4 Comparison of the tmtCETSA and diaCETSA**. **a** Workflows of tmtCETSA and diaCETSA. **b** Bar charts showing the number of total proteins, kinases, significant kinases, and significant non-kinases by using tmtCETSA and diaCETSA. **c** Pearson correlation coefficient of protein intensities between two replicates by using tmtCETSA and diaCETSA. **d** Violin plots showing the distributions of CVs of protein intensities between tmtCETSA and diaCETSA (n = 5 technical replicates). **e** Volcano plot visualization of kinase targets from K562 cell lysates, performed at 52 ℃ using 20 μM staurosporine by using tmtCETSA and diaCETSA. Adjust p-value=0.05 is indicated by a solid horizontal line. **f** Venn diagram displaying the kinase targets identified by classical TPP, tmtCETSA and diaCETSA methods. **g, h** Interaction map of significant proteins targets in the (**g**) tmtCETSA and (**h**) diaCETSA. (left panel) method, respectively. GO annotation of molecular function for all the identified significant proteins by using tmtCETSA and diaCETSA. Kinome tree displaying all staurosporine kinase targets identified by using tmtCETSA and diaCETSA. Circle size is proportional to the -log_10_ adjust p-value.

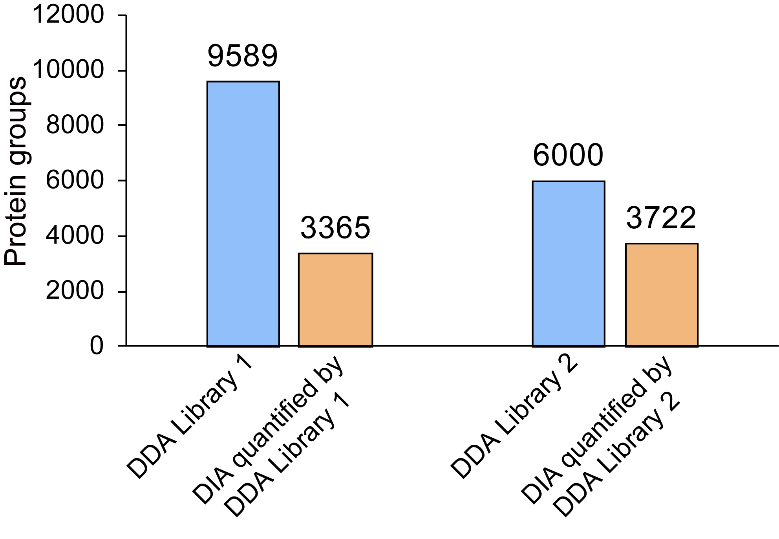

**Supplementary Fig. 5 The** **optimization of diaCETSA**. Bar charts showing DIA quantified protein groups by searching with different project-specific libraries.

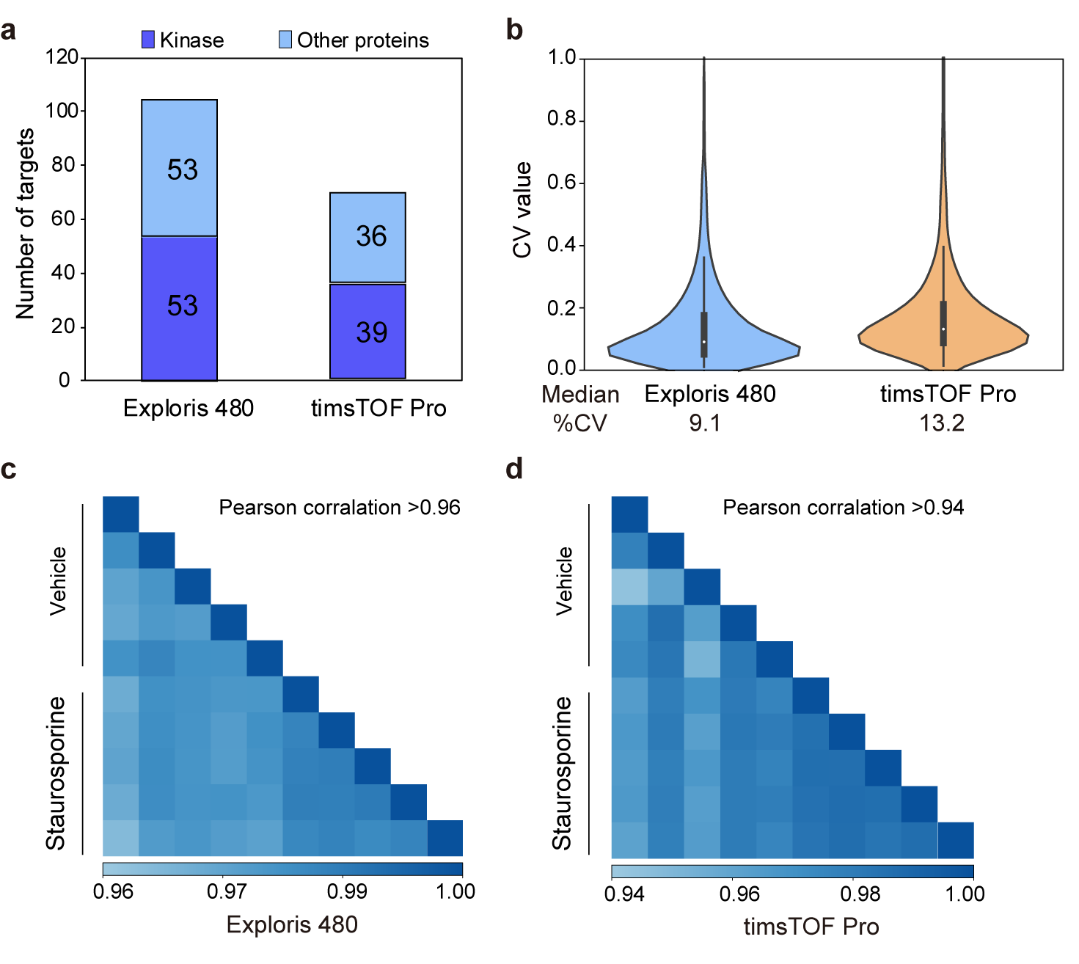

**Supplementary Fig. 6** **Comparison of the performance of Exploris 480 and timsTOF Pro for diaCETSA analysis**. **a** Histogram of the number of significant staurosporine targets. **b** Violin plots depicting CVs of protein intensities for the analysis of diaCETSA samples. **c, d** Pearson correlation coefficient of protein intensities for the diaCETSA samples in the (**c**) Exploris 480 and (**d**) timsTOF Pro experiments.

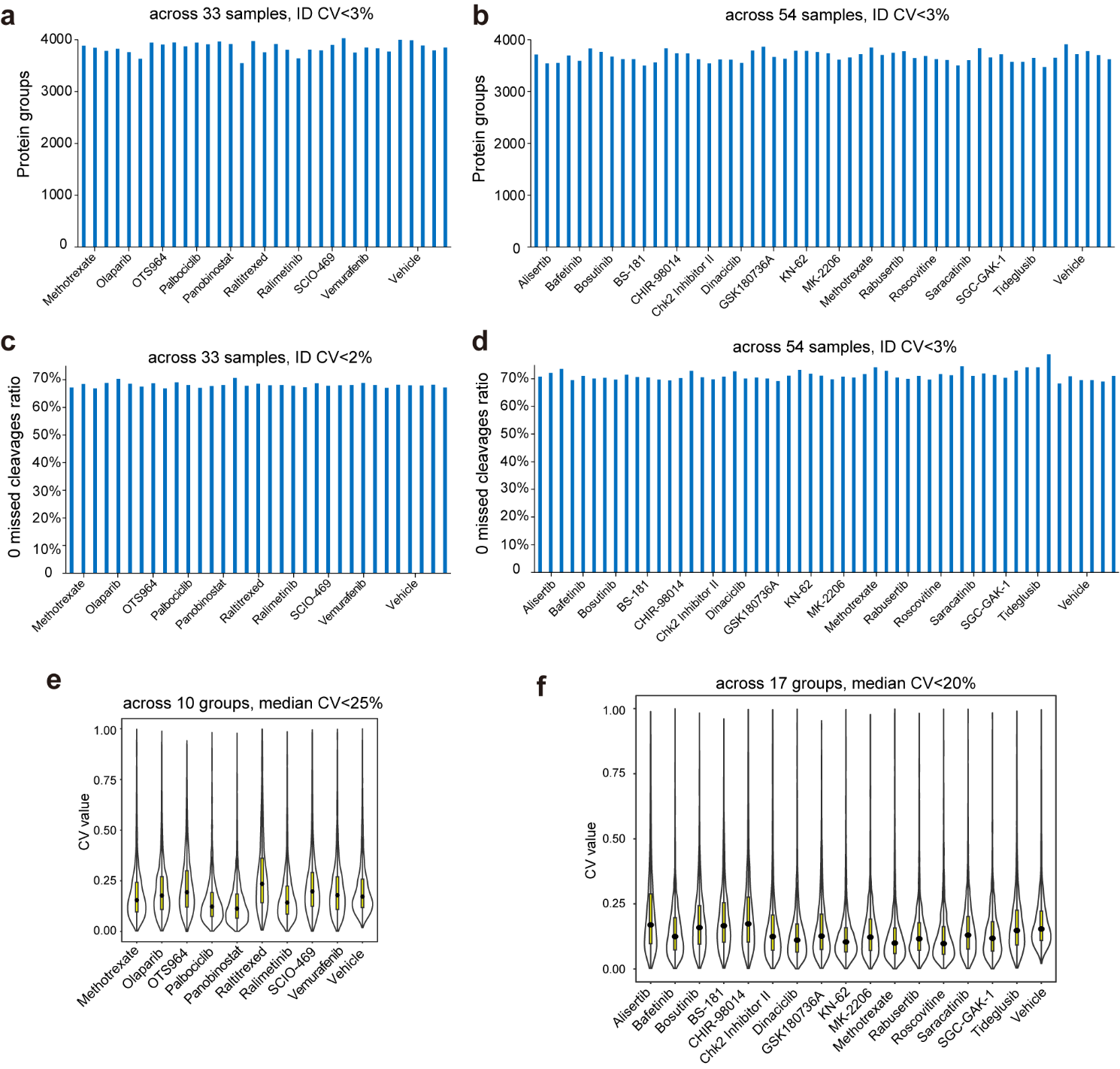

**Supplementary Fig. 7 Sample preparation performance of autoSISPROT for high-throughput drug target identification. a, b** The number of protein groups identified in **(a)** batch 1 (33 samples) and **(b)** batch 2 (54 samples). **c, d** The percentages of zero missed cleavages in **(c)** batch 1 and **(d)** batch 2. **e, f** Violin plots depicting CVs distribution of protein LFQ intensities from **(e)** batch 1 and **(f)** batch 2.
